# Euktect: Enhanced Eukaryotic Sequence Detection and Classification in Metagenomes via the DNA Language Model

**DOI:** 10.1101/2025.06.19.660294

**Authors:** Yibo Peng, Boyang Ji, Yinzhao Wang, Hongzhong Lu

## Abstract

The current taxonomy classification of DNA sequences in metagenomics primarily relies on alignment against reference databases. However, the eukaryotic species in genome databases are not sufficient as there exist numerous unculturable eukaryotes inside metagenomes. These limitations hinder functional and evolutionary analyses of eukaryotes across distinct environmental samples. To overcome these limitations, we created Euktect, a deep-learning-based toolbox for reliable, alignment-free classification of eukaryotic DNA sequences across different phylogenetic levels from metagenome datasets. Euktect achieves high accuracy in extraction of eukaryotic sequences longer than 500bp from the assembled contigs of metagenomes, significantly outperforming existing methods. Furthermore, we developed an algorithm that integrates Euktect’s predictions with existing tools to refine metagenome-assembled genomes (MAGs), substantially increasing the yield of high-quality eukaryotic MAGs for the downstream analysis. Beyond eukaryotic detection, Euktect incorporates two specialized models: a high-precision classifier for fungal phyla, and a hierarchical classifier that accurately identifies sequences from specific fungal genera (e.g., *Candida*) in diverse metagenomic samples. Significantly, this framework enables prediction of host disease status (e.g., inflammatory bowel disease) by linking eukaryotic sequence identification to clinical phenotypes through machine learning models. Collectively, Euktect enables accurate reconstruction and functional annotation of eukaryotic genomes from metagenomes at large scale, thereby empowering the comprehensive utilization of sequenced eukaryotic species for downstream evolutionary and clinical research.

## Introduction

Metagenomic research has historically focused on bacteria and archaea, but in fact, eukaryotic species are abundant in many metagenomic samples across various environments, including food^1^ oceans^2^, and soil^3^. Investigating eukaryotic species within metagenomes is essential for achieving a deeper understanding of microbial community composition, functionality, and evolutionary processes. Moreover, metagenomes contain a substantial number of uncultured, unidentified eukaryotic species. It is estimated that approximately 8.74 million eukaryotic species inhabit Earth^4^, yet the National Center for Biotechnology Information (NCBI) database contains only 18,631 eukaryotic reference genomes (before May 5, 2025). This gap indicates that the majority of eukaryotic species still lack comparable reference genomes in current databases,^5^, leading to a substantial difficulty for their identification and analysis. Thus, developing efficient methods to identify and characterize these unknown eukaryotic species from metagenomic data is a crucial and urgent task. The de novo construction of Metagenome-Assembled Genomes (MAGs) from metagenomes serves as an efficient approach for studying unknown environmental species. Nonetheless, the reconstruction of high-quality or medium-quality eukaryotic MAGs remains an unresolved challenge.

The identification of eukaryotic sequences in metagenomes has traditionally relied on alignment-based tools, including BLAST^6^, MMseqs2^7^, and Kraken2^8^. These tools assign the taxonomic label based on a genome similarity search of reference genome databases. The effectiveness of this comparison depends mainly on the quality and completeness of the reference databases used^9^. Since metagenomes may contain numerous unknown eukaryotic species absent from reference databases, alignment-based methods typically suffer from low recall and precision, and in some cases, fail entirely to identify such species. As a result, relying solely on alignment tools might not sufficiently facilitate the thorough extraction of eukaryotic sequences from metagenomic data.

To address the limitations of partial reference eukaryotic species databases, several alignment-free models based on machine learning and deep learning have been developed. Representative models include EukRep^10^, Tiara^11^, and BERTax^12^. EukRep is a linear Support Vector Machine (SVM) classifiers using sequence k-mer features. Tiara calculates term frequency–inverse document frequency (tf-idf) values based on k-mers, and utilize it to constructs a deep learning model. BERTax, based on the BERT (Bidirectional Encoder Representations from Transformers)^13^ language model, converts 3-mers in DNA sequences into tokens, transforming DNA into sentence-like structures for predictive analysis. These models enable the identification of eukaryotic sequences in metagenomes without requiring alignment to reference datasets. However, their accuracy for short sequences (1000 bp and below) remains limited, constraining their effectiveness in extracting shorter eukaryotic sequences.

In this study, we introduce Euktect, a deep learning model for taxonomic classification built upon the HyenaDNA^14^ architecture, a type of DNA language model capable of encoding DNA sequences at single-nucleotide resolution. We first constructed a classification model at the superkingdom level, enabling precise taxonomic classification of DNA sequences from eukaryotes, bacteria, archaea, and viruses. This model shows high predicted accuracy, particularly for sequences shorter than 1000 bp, outperforming current state-of-the-art model. We then applied this model to metagenomic analysis and developed an algorithm to refine eukaryotic MAGs. Our study demonstrate that the Euktect model effectively removes contaminating sequences from eukaryotic MAGs within metagenomes, significantly increasing the yield of high-quality and medium-quality eukaryotic MAGs. Furthermore, the model enables the direct extraction of eukaryotic sequences from metagenomic assemblies, assisting in downstream analyses of gene function and species composition. Extending beyond superkingdom-level classification, we developed a multi-class model for fungal phyla and a multi-level extraction model for the *Candida* genus, demonstrating the model’s considerable potential for eukaryotic species categorization and pathogenic fungus detection. The models presented here establish a robust computational framework for the identification and functional analysis of eukaryotic sequences in metagenomes, offering a novel perspective for exploring the interplay between species composition and functionality in metagenomic research.

## Results

### Construction of the Euktect model

Here, we introduce Euktect, an alignment-free taxonomic classification model built upon the HyenaDNA architecture. HyenaDNA^14^ is a Long-Range Genomic Foundation Model capable of encoding DNA sequences at single nucleotide resolution (Methods). In contrast to previous models like Eukrep^10^, Tiara^11^, and BERTax^12^, which utilize tokenizers or fixed k-mers, Euktect can capture a greater extent of the information inherent within a DNA sequence.

The workflow for constructing the superkingdom-level classification model is shown in Fig. 1a. To construct a dataset for superkingdom levels model training, we retrieved reference genomes from the NCBI database (released before April 14, 2024) using taxonomic IDs for each superkingdom, which included bacteria, archaea and microeukaryotes (fungi, protozoa and Chlorophyta). Due to the limited availability of viral reference genomes in NCBI, we downloaded all viral sequences, performed clustering to remove redundancy, and selected sequences longer than 5,000 bp to form the viral dataset. The resulting genomic sequences were shuffled and partitioned into training, validation, and test datasets in an 8:1:1 ratio. Using the HyenaDNA architecture, we employed a pretraining followed by fine-tuning approach (Methods). With the curated superkingdom dataset, we successfully constructed the Euktect model, which could be initially used for taxonomic classification at the superkingdom level. Two application examples of the model in metagenomics, including Eukaryotic MAG refinement and Eukaryotic sequence extraction as shown in Fig. 1b. Extension of the model in the fungal kingdom as illustrated in Fig. 1c.

**Figure 1.**
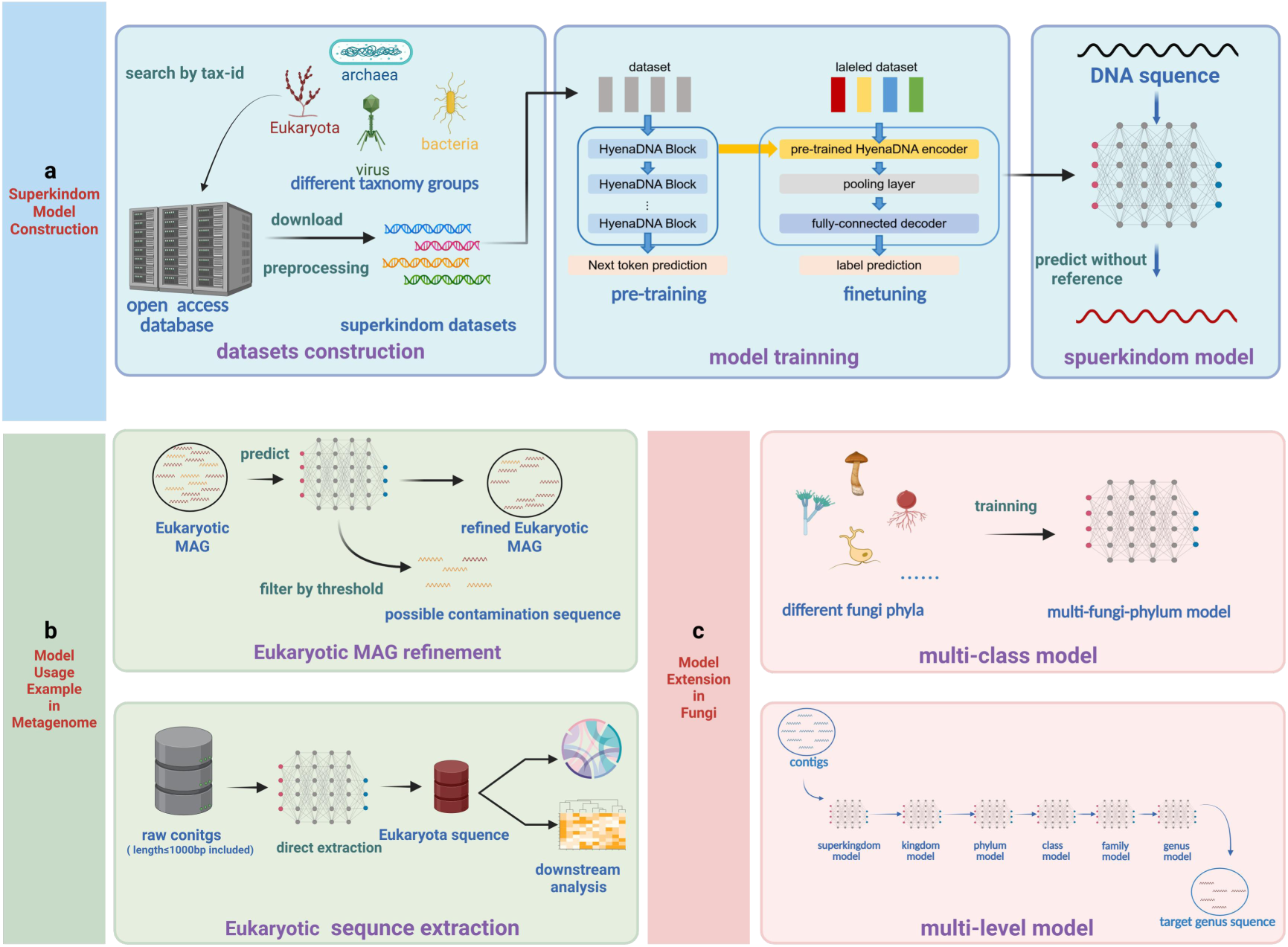
Overview of Euktect model. Euktect model construction pipeline at the superkingdom level. The construction of the Euktect model involves two main stages: dataset preparation and model training **(a)**. Two example applications of the Euktect model in metagenomic analysis, including refining eukaryotic Metagenome-Assembled Genomes (MAGs) and directly extracting eukaryotic sequences from metagenomic contigs for downstream analysis (**b**). Euktect model extension in the fungal kingdom. The Euktect model can be extended within the fungal kingdom to develop classification models for different fungal phyla. Additionally, it can be adapted into a hierarchical binary classification model to extract sequences from specific genera, enhancing its utility for targeted fungal genomic studies (**c**).

### Impact of hyperparameters and training method on the prediction performance of Euktect

We investigated the influence of hyperparameters and training method on the Euktect model’s performance, ultimately determining the optimal hyperparameters and training approach. Several adjustable hyperparameters can affect model performance. In this study, we used the superkingdom dataset to explore three key hyperparameters (Methods): model depth (layer number), model width (d_model), and model reverse bottleneck (d_inner). We evaluated validation loss across layer numbers of 1, 2, 3, 4, 5, and 6 (Fig. 2a). The results show that that increasing layer numbers greatly reduces validation loss, but as more layers are added, the change becomes less pronounced. Statistical analysis revealed significant differences in validation loss between 4 layers and 1–3 layers, but no significant differences between 4, 5, and 6 layers (Extended Data Fig. 1a). Next, we evaluated the impact of model width on predictive performance (Fig. 2b). Validation loss decreased markedly when increasing model width from 64 to 2048, and the prediction with model width at 2048 displayed significant differences compared to that at other values (Extended Data Fig. 1b). However, larger model widths increase parameter size and memory requirements, thus, in this work, we set the maximum model width at 2048 due to hardware constraints. Moreover, we assessed the performances of model when setting reverse bottleneck values at 128, 256, 512, 1024, and 2048 (Fig. 2c), respectively. It shows that the prediction accuracy of Euktect was only slightly improved (Extended Data Fig. 1c) when increasing reverse bottleneck value from 512 to 1014 or 2048. Based on these trials, we selected the following hyperparameters for superkingdom-level model training: model depth = 4, model width = 2048, and model reverse bottleneck = 512. Lastly, we examined how the training strategies influence model convergence profile during model training (Fig. 2d). It shows that the fine-tuned models from the pretrained one exhibited a significantly faster initial decline in validation loss compared to those trained from scratch (Extended Data Fig. 1d). This indicates that pretrained models incorporate prior knowledge of DNA sequence features, substantially enhancing the training efficiency.

**Figure 2.**
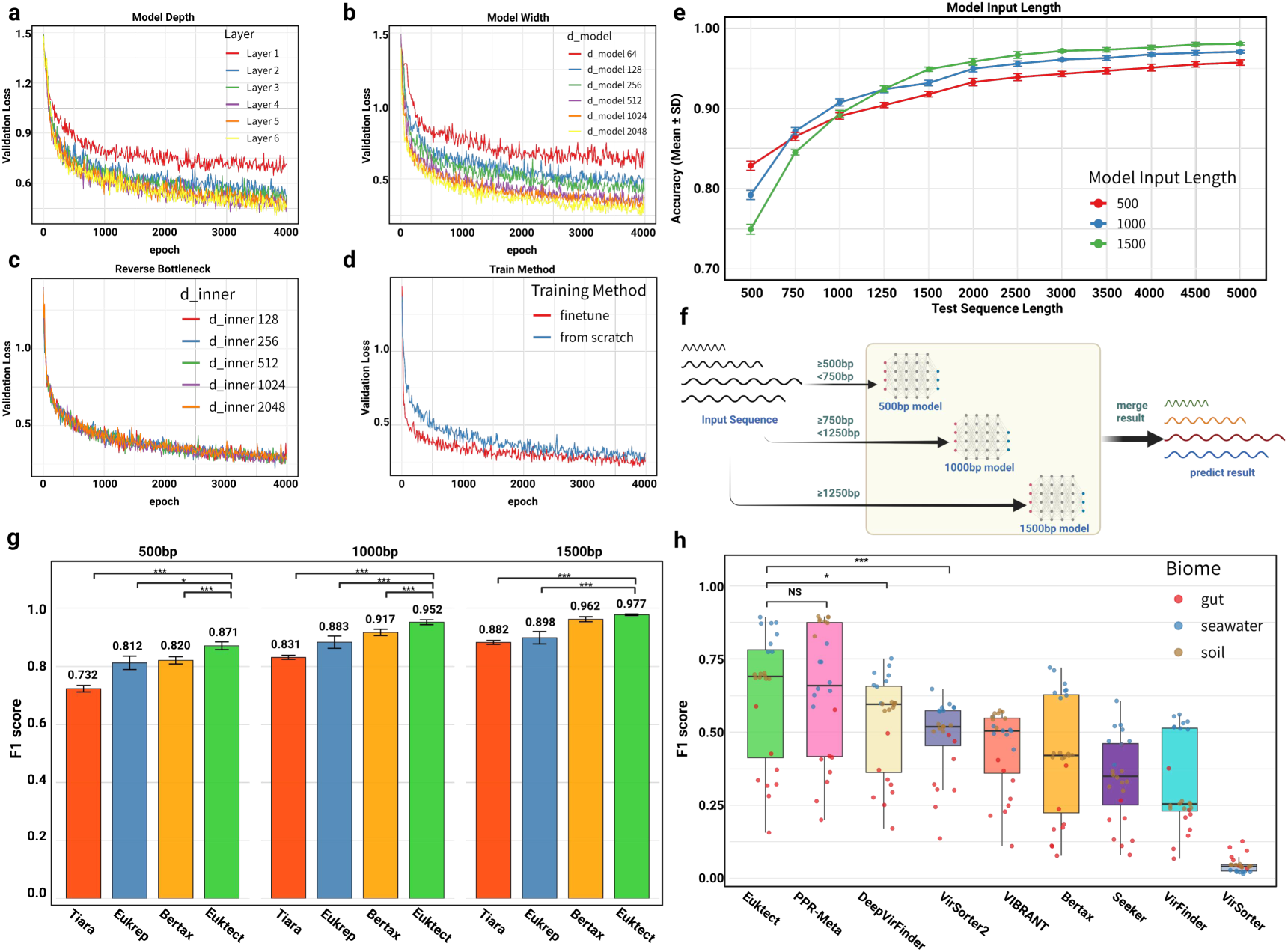
Impact of hyperparameters and training method on model performance and evaluation on benchmarks. Training loss under different model depths (**a**). Training loss under different model widths on training loss (**b**). Training loss under different reverse bottlenecks (**c**). Training loss under different training methods (**d**). The accuracy of models with different model input lengths under different test sequence lengths (**e**). Handling variable-length real sequence data with combined models (**f**). F1 Score comparison of Euktect and other tools on eukaryotic and bacterial identification benchmarks (**g**). F1 Score comparison of Euktect and other tools on viral identification benchmarks (**h**). ***: *P* value < 0.001, **: *P* value <0.01, *: *P* value < 0.05.

Within the assembled metagenome, the length of contigs could vary greatly, necessitating the training of classification models specifically designed for sequence with different length. We thus trained Euktect to generate different version of models, each of which has a fixed input sequence length. We then explored how the variance in the input sequence length influences the performance of those models (Methods). It shows that each model achieves optimal performance on sequences matching its training length: the 500 bp model for 500 bp sequences, the 1000 bp model for 1000 bp sequences, and the 1500 bp model for sequences of 1500 bp and longer. At a sequence length of 750 bp, the model with a 1000 bp input length slightly outperforms the model with a 500 bp input length. At a sequence length of 1250 bp, the performance of the models with 1500 bp and 1000 bp input lengths is generally comparable (Fig. 2e, Extended Data Figs 1e–f). By comparison, it indicates that the increased length of sequences for model training could enhance the prediction accuracy. To improve Euktect prediction performance for diverse lengths of DNA sequence, we used an ensemble technique that combined models trained with different input lengths (Fig. 2f). During applications, these sequences are assigned to the model that best fits their length range, resulting in optimal prediction performance for both short and long sequences.

### Euktect outperforms existing models

We tested Euktect against other alignment-free models and found that it outperforms the state-of-the-art models in classification of eukaryotic sequences. Given that eukaryotic and bacterial sequences predominate in most metagenomes, we first constructed a benchmark dataset for distinguishing eukaryotes from bacteria (Methods). We compared the performance of Euktect against EukRep^10^, Tiara^11^, and BERTax^12^ in classifying sequence at lengths of 500 bp, 1000 bp, and 1500 bp. For short sequences (500 bp and 1000 bp), Euktect demonstrated superior classification accuracy compared to other models (Fig. 2g). At 500 bp, Euktect achieved an F1 score of 0.871, representing a 5.1% improvement over BERTax’s score of 0.820. Similarly, at 1000 bp, its F1 score reached 0.952, exceeding BERTax by 3.5%. At 1500 bp, Euktect significantly outperformed Tiara and EukRep, achieving an F1 score of 0.977, a 1.5% improvement over BERTax (0.962).

Although Euktect primarily focuses on the prediction of eukaryotic sequences from metagenomes, it is, in fact, a superkingdom-level classification model capable of classify DNA sequences at the eukaryotic, bacterial, archaeal, and viral levels. To evaluate the broader applications of our model, the performance of the Euktect model in classifying virus sequences using benchmark datasets derived from real metagenomes^15^ was tested (Methods, Fig. 2h). The results show that our model’s performance in predicting viruses is comparable to PRP-Meta^16^, while it outperforms other virus prediction models^12,17–22^. Notably, PRP-Meta, the top-performing model for virus sequence identification, is specifically designed to classify sequences into viruses, chromosomes, and plasmids, but lacks the ability to further classify chromosome and plasmid sequences at taxonomic level. In contrast, Euktect model can not only identify sequences from viruses but also from eukaryotes, bacteria, and archaea, thus highlighting the versatility of our model in sequence taxonomic classification.

### Euktect contributes to reconstruction of more medium- and high-quality eukaryotic MAGs from metagenomes

De novo MAG construction from metagenomic data is crucial for functional studies of metagenomes. However, compared to prokaryotic MAG assembly, eukaryotic MAG construction faces substantial challenges. Eukaryotes are present in lower abundance when compared to prokaryotes in many metagenomic samples^23^, and their genomes are larger, often split across multiple chromosomes, with numerous non-coding and repetitive regions^24^. These factors significantly increase the complexity of eukaryotic genome assembly, making it challenging to construct medium-quality (MQ: completeness ≥ 50%, contamination ≤ 10%) and high-quality (HQ: completeness ≥ 90%, contamination ≤ 5%) eukaryotic MAGs according to the MIMAG^25^ standards originally defined for bacterial/archaeal MAGs. a challenging task. Furthermore, in metagenomic samples containing a mixture of eukaryotic, prokaryotic, and other species, genome binning tools such as MetaBAT2^26^ often produce bins with mixed eukaryotic and prokaryotic sequences, thus lowing the quality of MAGs and increasing the number of low-quality (LQ) eukaryotic MAGs accordingly. We conducted a binning simulation (Methods, Extended Data Figs 2). It was observed that as the proportion of bacterial sequences in the simulated contigs increased, the number of eukaryotic sequences in the eukaryotic bins generated by Metabat2 remained largely constant. However, there was a trend of increasing contamination by bacterial sequences.

To address the above issue, we developed an algorithm integrating the prediction from Euktect model with MAG quality assessment tools to refine eukaryotic MAGs, and the result shows that this algorithm significantly increases the number of medium-quality (MQ) and high-quality (HQ) eukaryotic MAGs assembled from metagenome datasets (Fig. 3a). The algorithm begins by using the Euktect model to predict the eukaryotic origin probability for each sequence within a MAG obtained from binning tools. A user-defined threshold is then applied to filter out sequences with eukaryotic probabilities below this value. The filtered MAG is subsequently evaluated using a quality assessment tool, such as EukCC^27^, to calculate completeness and contamination, which will be used to classify the refined MAG into three groups – HQ, MQ and LQ. See detailed algorithm description in (Methods).

**Figure 3.**
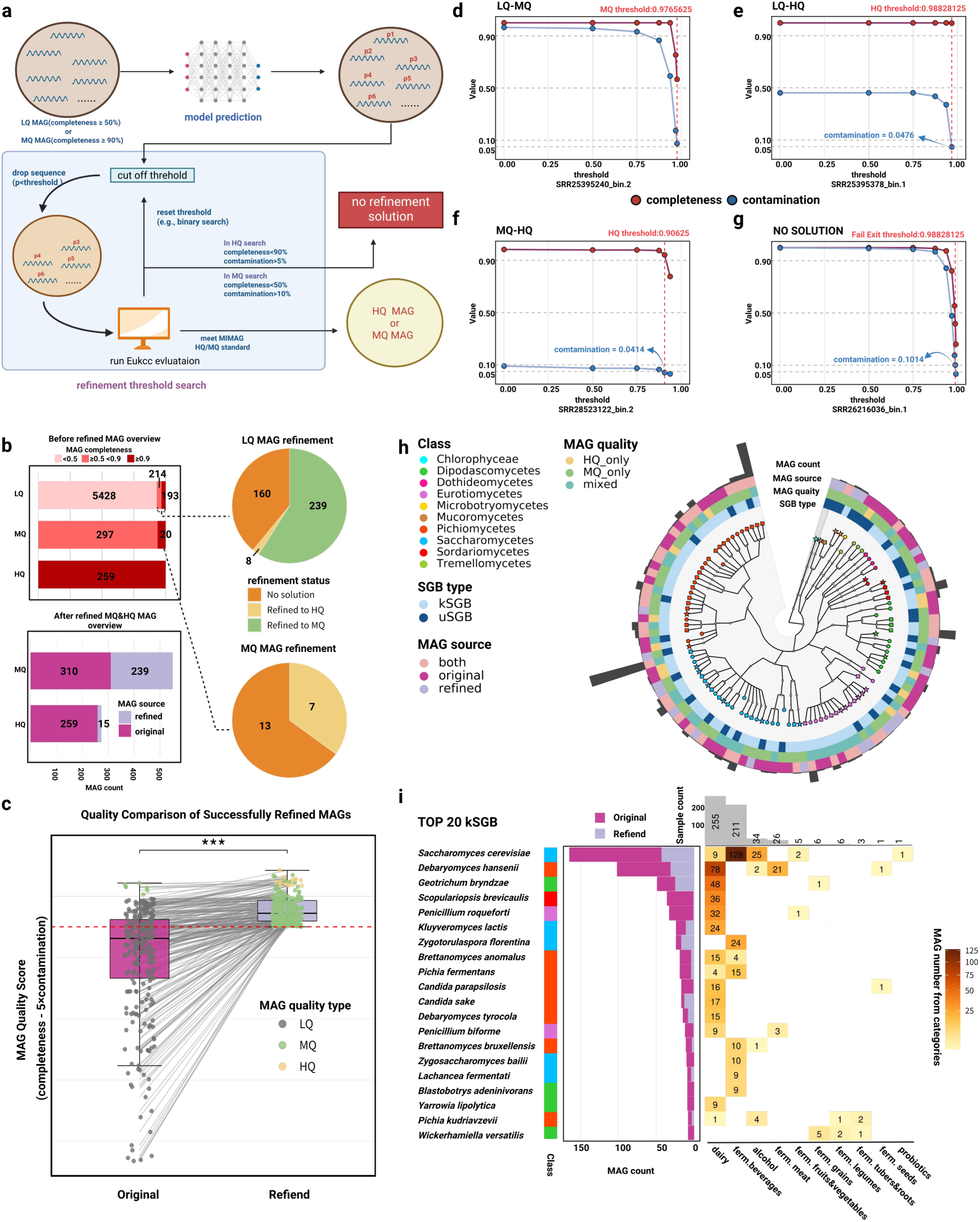
Euktect usage for eukaryotic MAGs refinement algorithm. Eukaryotic MAGs refinement algorithm scheme (**a**). Overview of refinement algorithm performance on real metagenomic datasets (**b**). Comparison of successfully refined MAGs quality score (completeness −5 × contamination) before and after refinement. (**c**). Changes in completeness and contamination during LQ to MQ MAG refinement (**d**). Changes in completeness and contamination during LQ to HQ MAG refinement (**e**). Changes in completeness and contamination during MQ to HQ MAG refinement (**f**). Changes in completeness and contamination during No Solution MAG refinement (**g**). SGB clustering and species annotation of MQ and HQ eukaryotic MAGs (**h**). Distribution of Top 20 kSGBs across different source categories (**i**).

Our MAG refinement algorithm could significantly increase the number of MQ and HQ MAGs reconstructed from metagenomes. We applied this algorithm to a large-scale eukaryotic MAG construction and refinement process on food metagenome dataset^1^ (Methods). The refinement results are shown in Fig. 3b. Initially, the MAGs constructed from dataset contained 407 LQ MAGs with completeness ≥ 50% and 20 MQ MAGs with completeness ≥ 90%. After refinement, 239 MQ MAGs and 8 HQ MAGs were obtained from the LQ MAGs, while 7 HQ MAGs were derived from the MQ MAGs. Ultimately, the dataset yielded 310 MQ MAGs (including 239 refined) and 259 HQ MAGs (including 15 refined). The MAG quality score (QS), defined as completeness − 5 × contamination^28^, was analyzed before refinement for LQ and MQ MAGs (Extended Data Figs 3a and 3b). LQ-to-HQ refined MAGs exhibited significantly higher QS than LQ-to-MQ MAGs, which in turn had higher QS than LQ MAGs with no solution. Similarly, MQ-to-HQ refined MAGs had significantly higher QS than MQ MAGs with no solution, indicating that refinement success is strongly correlated with initial MAG quality. Post-refinement QS comparisons (Fig. 3c) revealed significant improvements in QS for successfully refined MAGs.

To illustrate potential outcomes during the refinement process, we present four representative eukaryotic MAGs as example. Fig. 3d illustrates the refinement of a LQ MAG into an MQ MAG, showing trends in completeness and contamination as the filtering threshold increases. Initially, the LQ MAG exhibited 100% completeness along with a high contamination rate of 95.65%. As the threshold increased, both completeness and contamination decreased, with contamination declining more rapidly. At a threshold of 0.9765625, the MAG achieved a completeness of 56.73% and contamination of 7.45%, meeting the MQ standard. Fig. 3e depicts the refinement of an LQ MAG into an HQ MAG, where completeness decreased slightly from 99.79% to 99.59%, while contamination dropped significantly from 46.17% to 4.76%, meeting the HQ standard. Fig. 3f shows the refinement of an MQ MAG into an HQ MAG. In contrast, Fig. 3g illustrates a “no solution” case, where, at a threshold of 0.98828125, the completeness of the MAG was 41.82% and its contamination remained at 10.14%, failing to meet the MQ standard even after the filtering process using our algorithm. The relationship between the threshold values for successful MAG refinement and the QS before refinement is illustrated in (Extended Data Figure 3c). Since the refinement algorithm operates through sequence filtering, a significant limitation of this approach is the reduction in MAG size. The correlation between the size reduction ratio of successfully refined MAGs and the MAG QS before refinement is presented in (Extended Data Figure 3d). Notably, MAGs with initially lower QS values exhibit an increasing trend in size reduction ratio, which may be attributed to the potentially higher proportion of contaminating sequences in these MAGs.

We performed Single Genome Bin (SGB) clustering and taxonomic labeling on all resulting MQ and HQ MAGs (Methods). The results (Fig. 3h) revealed that some SGBs were derived exclusively from refined MAGs, many of which were unknown SGBs (uSGBs). This suggests that the refinement algorithm enables the reconstruction of genomes from previously uncharacterized species in metagenomic samples. Additionally, we analyzed the distribution of the top 20 known SGBs (kSGBs) across samples (Fig. 3i). The most abundant kSGB was Saccharomyces cerevisiae, with 122 original MAGs and an additional 43 MAGs obtained through refinement. By generating more MQ and HQ MAGs for known species, the refinement algorithm facilitates systematic studies of their distribution and evolutionary patterns across diverse samples.

### Euktect enables isolation of eukaryotic sequences from metagenomes

Traditional metagenomic studies often focus on bacteria, archaea, or entire microbial communities. For studies targeting eukaryotic sequences, isolating these sequences from mixed samples is a prerequisite. Owing to the high prediction accuracy of Euktect, it could be used to directly isolate eukaryotic sequences from metagenomic samples, thus enabling targeted downstream analyses of eukaryotic components. In this regard, we reanalyzed 173 metagenomes from various food samples^1^. All sequenced reads were preprocessed and assembled into contigs (Methods), and eukaryotic sequences were extracted using the Euktect model. While previous metagenomic studies typically discarded contigs shorter than 1000 bp^1^, our prior evaluation showed that Euktect has high accuracy even for classifying 500 bp sequences, allowing us to extend eukaryotic sequence filtering to contigs ≥ 500 bp.

In the food metagenome, which exhibited high eukaryotic sequence abundance, we analyzed the extracted eukaryotic sequences. A straightforward downstream application is estimating eukaryotic sequence abundance by mapping extracted sequences back to the original reads and calculating relative abundance (Methods). The estimated eukaryotic abundance across 173 food metagenome samples, categorized by sample type, is shown in Fig. 4a. Cheese rind samples showed a significant increase in eukaryotic sequence abundance post-ripening compared to pre-ripening, while sausage and whey samples exhibited significant decreases post-ripening. Cheese core, cured meat, and fresh meat samples showed no significant changes in eukaryotic abundance before and after ripening.

**Figure 4.**
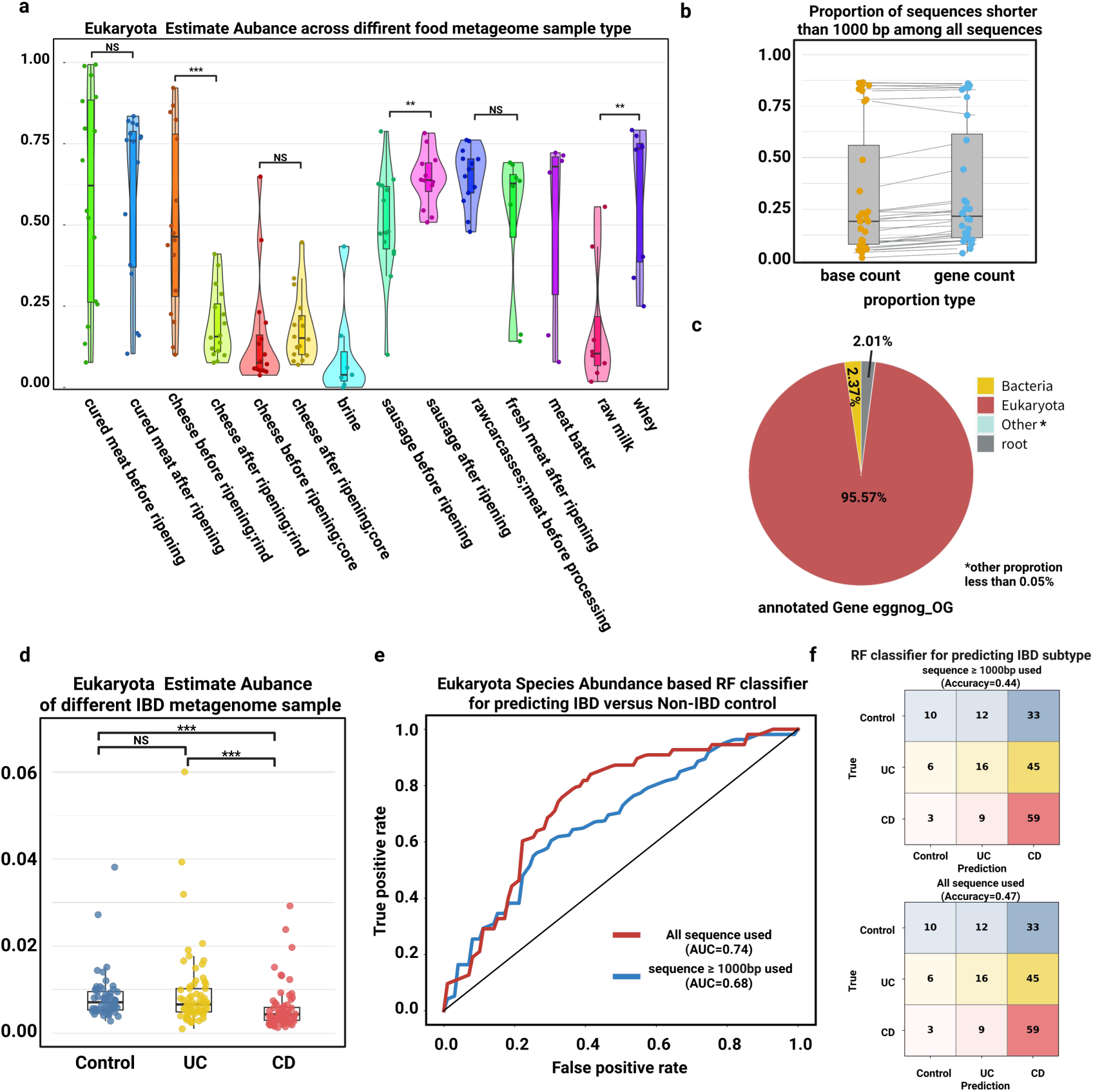
Euktect usage for direct extraction of eukaryotic sequences from metagenomes for downstream analysis. Abundance estimation of different eukaryotic sequence categories in Food metagenomic samples. The Euktect model was used to estimate the abundance of various categories of eukaryotic sequences within food metagenomic samples (PRJNA997800) (**a**). For the cheese-type metagenome samples, downstream gene annotation was performed. Subsequently, for samples containing more than 10,000 genes, the number of annotated genes was counted. Two key proportions were also determined per sample: the ratio of base pairs from sequences shorter than 1,000 bp to the total, and the ratio of annotated genes on those short sequences to the total. (**b**). Proportion of different eggNOG orthologous groups in all annotated genes (**c**). Abundance estimation of eukaryotic sequences in IBD-Related metagenomic samples (**d**). ROC curve for non-IBD vs. IBD prediction using random forest model based on eukaryotic species abundance (**e**). Confusion matrix for IBD subtype prediction using random forest model based on eukaryotic species abundance (**f**). ***: *P* value < 0.001, **: *P* value <0.01, *: *P* value < 0.05.

We performed gene-level analysis on eukaryotic sequences extracted from cheese core (before and after ripening) and cheese rind (before and after ripening) samples. Low-abundance samples (< 0.10) were filtered out, and gene annotation was conducted using eggNOG-mapper^30^. Notably, a substantial number of genes were annotated even in sequences shorter than 1000 bp. The proportion of bases from 1000 bp sequences and the proportion of genes predicted from these sequences relative to the total are shown in Fig. 4b. These proportions were highly correlated (Pearson correlation = 0.9969). In samples with a high proportion of bases from sequences < 1000 bp, the proportion of predicted genes from these sequences was also high. This is partly due to reduced specificity in gene prediction for shorter sequences^31^, leading to more false positives. However, given that sequences < 1000 bp often dominate metagenomic contigs^32^, these sequences may contain valuable genes warranting further investigation, even after accounting for overprediction.

For annotated genes, we analyzed their eggNOG Orthologous Groups (OGs) (Fig. 4c), finding that 95.57% of annotated genes were eukaryotic, indirectly validating the reliability of our extracted eukaryotic sequences. Gene abundances were calculated and regrouped by KEGG Ortholog (KO)^33^. KEGG pathway enrichment analyses were performed for genes from cheese rind and cheese core samples (before and after ripening) (Methods). Enrichment results for cheese rind (Extended Data Fig. 4a) showed a greater number of enriched pathways post-ripening, reflecting the significant increase in eukaryotic sequence abundance. In contrast, cheese core samples showed fewer enriched pathways post-ripening (Extended Data Fig. 4b), consistent with their minimal change in eukaryotic abundance. Enriched pathways in cheese core samples were primarily associated with amino acid, cysteine, methionine, galactose, and selenocompound metabolism, linked to flavor and nutritional properties.

### Eukaryotic sequences extracted by Euktect facilitated the disease prediction

Eukaryotic sequences within metagenome could be used as the biomarkers to predict the host disease conditions. Here, using a similar procedure, we reanalyzed 220 metagenomes related to inflammatory bowel disease (IBD)^29^. After quality control analysis, 193 high-quality samples were retained for downstream analysis (Methods). Euktect was used to extract eukaryotic sequences. IBD comprises two main subtypes: ulcerative colitis (UC) and Crohn’s disease (CD). Eukaryotic sequence abundances across non-IBD control, UC, and CD samples are shown in Fig. 4d. No significant differences were observed between control and UC samples, but CD samples exhibited significantly different eukaryotic abundances compared to both control and UC samples. Species annotation was performed on the extracted sequences, and eukaryotic species abundances were calculated using all sequences and those ≥ 1000 bp. A random forest (RF) binary classification model was built to distinguish non-IBD from IBD samples based on eukaryotic species abundance, with fivefold cross-validation (Methods). The receiver operating characteristic (ROC) curves (Fig. 4e) show that the model using all extracted sequences achieved an area under the curve (AUC) of 0.74, 6% higher than the model only using sequences ≥ 1000 bp (AUC = 0.68), suggesting that including all sequences provides a more accurate representation of species abundance. Additionally, an RF model for predicting IBD subtypes was constructed with fivefold cross-validation (Fig. 4f). The model using all sequences achieved an accuracy of 0.47, compared to 0.44 for the model using sequences ≥ 1000 bp.

### Extension of the Euktect model to fungal phyla for multi-classification

Classifying fungi based on sequence has long been a critical challenge in fungal biology. Traditionally, fungal classification relies heavily on nuclear internal transcribed spacer (ITS) sequences^34^. However, most sequences in metagenomic datasets lack ITS regions, limiting the utility of such approaches. To address this issue, we extended the superkingdom-level Euktect model to sub-models, which could be used to classify sequences across 10 distinct fungal phyla using the same training method (Methods). The performance of these models was evaluated on a test dataset, with confusion matrices for 1000 bp sequences shown in Fig. 5a and F1 scores across different sequence lengths presented in Fig. 5b. The results reveal varying prediction accuracies across fungal phyla. To investigate this variability, we analyzed the relationship between the number of reference genomes in the training dataset and the F1 score for each phylum (Extended Data Fig. 5). It was observed that as the number of reference genomes increases, the prediction accuracy of the model exhibits a declining trend. This may be attributed to the increased sequence diversity within the phylum as more reference genomes are included, which consequently reduces the model’s prediction accuracy for this phylum.

**Figure 5.**
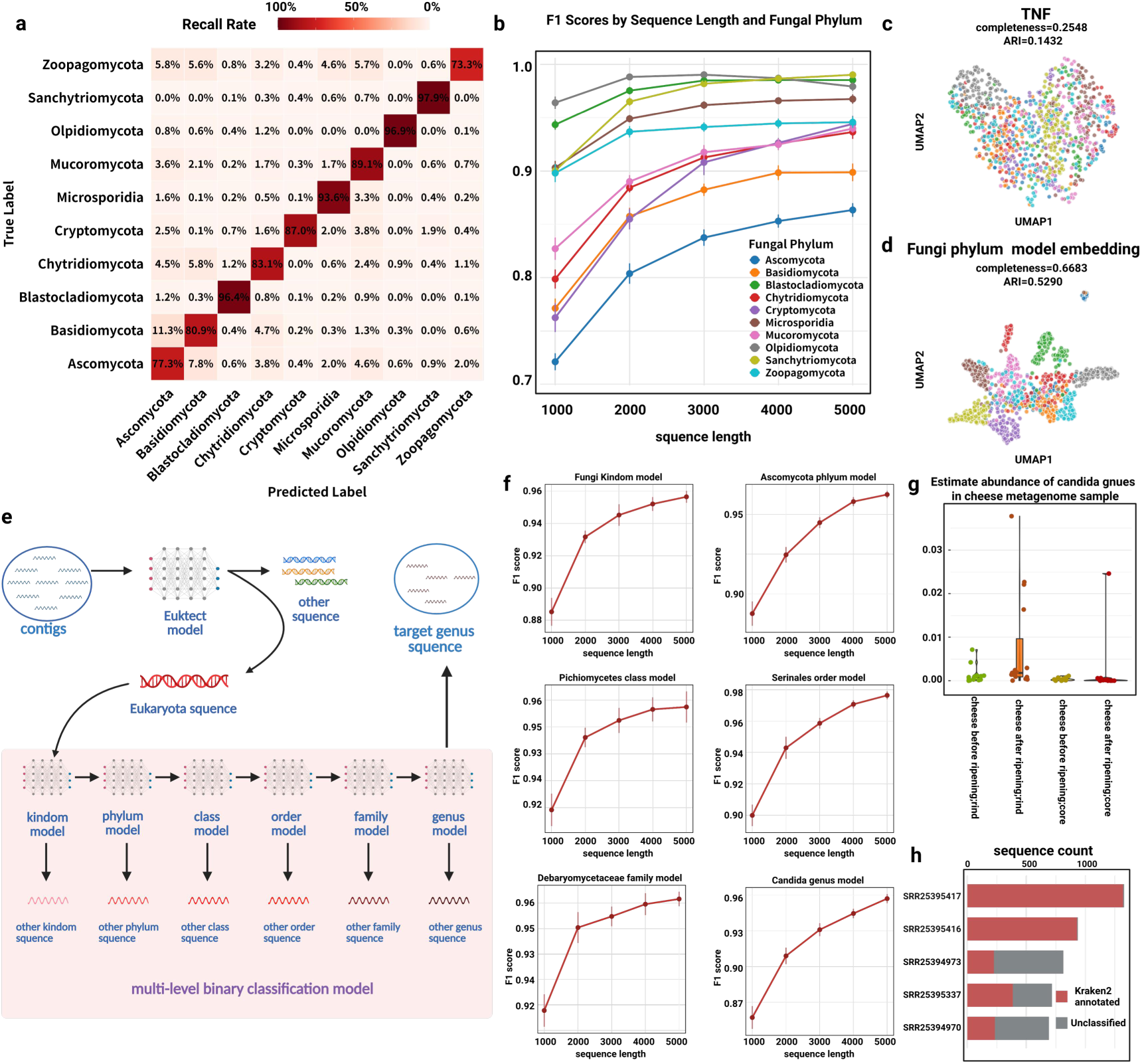
Model extension to the fungal kingdom. Confusion matrix of the fungal phyla multi-classification model on the test sequence at 1000 bp length (**a**). F1 score variation of the fungal phyla multi-classification model across test sequence length (**b**). Clustering results of fungal sequences from different phyla using TNF embedding. Sequences from different fungal phyla were embedded using the Tetranucleotide Frequency (TNF) method, followed by clustering with the K-means algorithm (**c**). Clustering results of fungal sequences using the encoder of the fungal multi-classification model. The encoder of the fungal multi-classification model was used to embed sequences from different phyla, followed by clustering with the K-means algorithm (**d**). *Candida* Genus multi-level model scheme. The hierarchical model begins with the Euktect model extracting eukaryotic sequences from contigs. For sequences targeting specific genera, direct genus-level multi-classification may lack sufficient accuracy. Instead, a hierarchical binary classification model is employed to extract target genus sequences with improved accuracy while maintaining robustness. Using the *Candida* genus as a case study, a multi-level extraction model was developed, encompassing models for the Fungi kingdom, *Ascomycota* phylum, *Pichiomycetes* class, *Serinales* order, *Debaryomycetaceae* family, and *Candida* genus (**e**). F1 Score variation across test sequence lengths for multi-level models (**f**). Abundance estimation of *Candida* genus in cheese samples (**g**). Annotation of *Candida* genus sequences in top five samples using reference genomes. For the five samples with the highest number of extracted *Candida* genus sequences, annotation was performed using reference genomes (**h**).

We further explored sequence clustering for the 10 fungal phyla (Methods). Clustering based on Tri-Nucleotide Frequency (TNF) is shown in Fig. 5c, while clustering based on embeddings generated by our model is shown in Fig. 5d. The latter approach successfully separated sequences at the phylum level, particularly for phyla with fewer reference genomes, such as *Olpidiomycota*, *Sanchytriomycota*, *Blastocladiomycota*, and *Microsporidia*, which were distinctly separated from other phyla. In contrast, larger phyla like *Ascomycota* and *Basidiomycota* exhibited great overlap with other phyla, making them harder to distinguish. This may be attributed to the reliance of these phyla’s classification on morphological traits rather than genomic sequence similarity.

### Multi-Level model for effective extraction of *Candida* genus sequences

*Candida* species are major components of fungal communities in various human body sites, with several species, such as *Candida albicans* and *Candida tropicalis*, being pathogenic^35^. Identifying *Candida* species in metagenomic samples is highly valuable, but the limited number of *Candida* reference genomes (33 as of May 5, 2025) and the potential presence of unknown *Candida* species pose challenges for accurate taxonomic analysis of *Candida* species. Direct genus-level classification models often yield low accuracy; for instance, the BERTax model reported a macro-average precision of only 66.92% at the genus level. To overcome this issue, we constructed a multi-level binary classification model to extract *Candida* sequences from metagenomic samples (Methods, Fig. 5e). This model encompasses binary classifiers at the fungal kingdom, Ascomycota phylum, Pichiomycetes class, Serinales order, Debaryomycetaceae family, and *Candida* genus levels, maintaining high prediction accuracy at each level with F1 scores exceeding 95% for 5000 bp sequences (Fig. 5f).

We applied this multi-level model to extract *Candida* sequences from four types of metagenomes from the above food samples^1^: cheese core before ripening, cheese core after ripening, cheese rind before ripening, and cheese rind after ripening. Estimated *Candida* abundance across these environments is shown in Fig. 5g. Extracted sequences were annotated using the alignment-based tool Kraken2 against *Candida* reference genomes. The top five annotated sequences are presented in Fig. 5h. Sequences from samples SRR25395417 and SRR25395416 (both cheese rind before ripening) showed strong alignment with reference genomes, predominantly annotated as *Candida tropicalis*. In contrast, sequences from samples SRR25394973, SRR25395337, and SRR25394970 exhibited lower proportions of matches to reference genomes. These unclassified sequences may result from model prediction errors or, more likely, represent sequences from unknown *Candida* species absent from reference databases.

## Discussion

In recent years, advancements in metagenomics and associated technologies have enabled the sequencing of an increasing number of metagenomic samples, offering deeper insights into the biological diversity of natural environments. However, current reference databases encompass genomic data for only a small fraction of Earth’s species, leaving the vast majority uncharacterized. While most metagenomic research has focused on bacteria and archaea, microeukaryotes in the environment have received relatively little attention. Traditional sequence alignment methods are often insufficient for identifying unknown species and eukaryotic sequences within metagenomes. To overcome the limitations imposed by alignment on existing databases for uncovering unknown species, deep learning models based on k-mer and BERT architectures have been proposed. These models enable alignment-free identification of sequences, aiding in the elucidation of their biological classification. Building on recent developments in large language models, we developed the Euktect model using the HyenaDNA^14^ architecture, which allows for precise identification of DNA sequence origins at the superkingdom level. A key advantage of Euktect is its ability to achieve higher prediction accuracy on shorter sequence lengths compared to previous models. In conjunction with the Euktect model, we introduced a novel algorithm for refining eukaryotic MAGs, facilitating the reconstruction of more medium-quality and high-quality eukaryotic MAGs from metagenomes. Moreover, we demonstrated the model’s scalability across biological classification levels, including multi-class classification at the fungal phylum level and multi-level models targeting specific genus. The suite of models presented in this study provides reliable tools for diverse tasks in metagenomic analysis, particularly addressing the critical gap in efficient methods for analyzing eukaryotic organisms within metagenomes.

Despite these innovations, several limitations persist. First, the availability of high-quality genomic data for training remains constrained, with current reference genomes estimated to represent only about 2% of all eukaryotic species. Although our model performs robustly at the fungal phylum level—achieving an average F1 score above 80% for sequences as short as 1000 bp—its predictive accuracy declines when applied directly to finer taxonomic resolutions, such as class or genus levels. As a result, constructing multi-level binary classification models is necessary to achieve relatively accurate identification for specific genus. Additionally, the HyenaDNA architecture, being a relatively new framework, lacks extensive research on its interpretability, rendering the model a “black box” and complicating efforts to analyze its decision-making processes.

The ongoing advancement of deep learning model architectures, training approaches, and datasets offers significant potential for enhancing sequence prediction accuracy. Extending the biological classification capabilities of the model to the genus level, thereby achieving alignment-free multi-classification, could greatly advance diversity studies at finer taxonomic resolutions within metagenomic samples. Furthermore, conducting interpretability analyses on the model could provide deeper insights into the molecular mechanisms and sequence features underlying species classification. Such progress may enable the identification of differentiation patterns among bacteria, archaea, and eukaryotes, potentially offering clues to the origins and evolutionary trajectories of life.

## Methods

### Dataset preparation

The datasets used in this study were constructed using NCBI command-line tools^36^ to retrieve genomic data from the NCBI database, with biological classification labels assigned according to the NCBI taxonomy database^37^. For training the Euktect model, we collected reference genomes from NCBI (release before April 14, 2024), including 5,277 microeukaryote reference genomes, 18,650 bacterial reference genomes, and 677 archaeal reference genomes. Due to the limited availability of viral reference genomes, we downloaded all available viral sequences and performed clustering using MMseqs2^7^ (with parameters set to --min-seq-id 0.9 --cov-mode 0 -c 0.8) to remove redundant sequences. Sequences with lengths ≥ 5,000 bp were retained to form the viral dataset. All downloaded data are stored in dehydrated form in the Supplementary Dataset 1 and can be rehydrated using NCBI command-line tools to obtain the complete genomic dataset. The downloaded sequences for eukaryotes, bacteria, archaea, and viruses were randomly shuffled within each group and partitioned into training, validation, and test datasets in an 8:1:1 ratio. This process generated the dataset used for constructing the superkingdom-level classification model.

### Model architecture

The Euktect model is built upon the HyenaDNA architecture^14^, a genomic foundation model composed of stacked Hyena blocks. Each Hyena block primarily consists of Hyena operator^38^ and Feed-Forward Network (FFN). The Hyena operator employs a parameter-efficient global convolutional filter combined with a data-controlled gating mechanism. For an order-N Hyena operator, the output (y) can be expressed as:

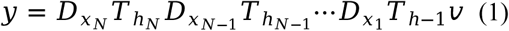

Where *v* ∈ ℝ*^L^* is the initial projection of the input sequence, *x_n_* ∈ ℝ*^L^* is the gating projection at step n, 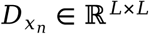 is a diagonal matrix with x_n_ on its diagonal, 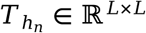 is a lower-triangular Toeplitz matrix representing the causal convolution with filter h_n._

*v* ∈ ℝ*^L^* Compared to the BERT architecture, the HyenaDNA architecture can handle longer token sequences. For DNA sequence tokenization, it treats each nucleotide as an individual token, enabling single-nucleotide resolution encoding. The token types include “A”, “G”, “C”, “T”, and “N”, along with additional special tokens for padding, separation, and unknown characters.

### Model hyperparameters

The Euktect model’s primary hyperparameters are as follows:

1. Model Depth (Layer Number): The depth of the model, corresponding to the number of stacked Hyena blocks.
2. Model Width (d_model): The width of the model, defined as the dimension of the hidden state in each Hyena block.
3. Model Reverse Bottleneck (d_inner): Also referred to as the FFN Expansion Factor, this parameter specifies the dimension of the intermediate layer in the FFN within each Hyena block.
4. Input Length: The model requires a fixed input sequence length. During prediction, sequences shorter than the input length are padded with “N” characters to match the input length. For input sequences exceeding the fixed input length, the sequence is divided into consecutive non-overlapping chunks of equal length. If the total number of chunks does not exceed a predefined max chunk number, an additional chunk is appended by extracting the final segment of the original sequence, equal in length to the model input, thereby potentially overlapping with the previous chunk. This ensures the inclusion of terminal sequence information that might otherwise be omitted. Each chunk is processed independently, and the final prediction for the entire sequence is obtained by averaging the prediction scores across all processed chunks.

### Model training

The Euktect model was trained using two approaches: direct supervised learning from scratch and a pretraining followed by fine-tuning strategy. In the supervised learning approach from scratch, labeled datasets were input into the initial Euktect model to perform sequence label prediction, resulting in a model for taxonomic classification. The second approach involved pretraining and fine-tuning. During pretraining, the model was trained on unlabeled data using a next-token prediction task, where the model predicts the next nucleotide in a sequence based on the preceding context. The loss function for pretraining is defined as:

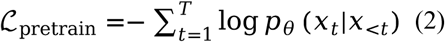

Where x_t_ is the true nucleotide at position t. *p_θ_*(*x_t_*|*x*_<<u>t</u>_) is the predicted probability of nucleotide x_t_ given the preceding context.

*p_θ_*(*x_t_*|*x*_<<u>t</u>_) This pretraining phase enables the model to learn inherent patterns in DNA sequences. Following pretraining, the model underwent fine-tuning on a labeled dataset for sequence label prediction. The loss function for fine-tuning is defined as:

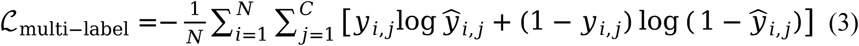

Where N = number of samples, C = number of labels, *y_i,j_* ∈ {0,1}is the ground-truth binary label for sample i and class j, 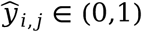 is the predicted probability of class j being active in sample i.

The fine-tuned model is capable of predicting labels for DNA sequences, corresponding to their taxonomic classification. The code used for model training is available in the HyenaDNA GitHub repository: https://github.com/HazyResearch/hyena-dna. Training was conducted on a single NVIDIA A100 40GB GPU for three input lengths (500 bp, 1000 bp, and 1500 bp) with the following hyperparameters: learning rate (lr) = 6e-5, model width (d_model) = 2048, number of layers (n_layer) = 4, model reverse bottleneck (d_inner) = 512, and maximum epochs = 10,000. Pretrained models were first constructed for each input length and subsequently fine-tuned to obtain the final models.

### Model evaluation

#### Evaluation across different sequence lengths

To assess model performance across various sequence lengths, test datasets were constructed as follows. From the four test datasets (eukaryotes, bacteria, archaea, and viruses) generated during the dataset preparation phase (see Dataset Preparation), 1,000 sequences of 500 bp were randomly sampled from each group, resulting in a total of 4,000 sequences. This process was repeated 10 times to create 10 test datasets for evaluation at 500 bp. Similarly, test datasets were constructed for sequence lengths of 750 bp, 1000 bp, 1250 bp, 1500 bp, 2000 bp, 2500 bp, 3000 bp, 3500 bp, 4000 bp, 4500 bp, and 5000 bp. Models with input lengths of 500 bp, 1000 bp, and 1500 bp were evaluated on these datasets using prediction accuracy as the evaluation metric, defined as:

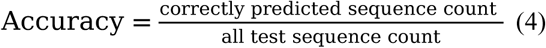

Prediction results are provided in the Supplementary Dataset Prediction Across Length Result.

#### Eukaryotic identification benchmark datasets

To ensure an objective evaluation of eukaryotic prediction tools and avoid data leakage, we downloaded eukaryotic and bacterial reference genomes released between April 14, 2024, and November 17, 2024, using NCBI command-line tools. These genomes were not included in the training datasets of any evaluated tools. Following a similar process to the test dataset construction, 1,000 sequences of 500 bp were randomly sampled from both eukaryotic and bacterial reference genomes, resulting in 2,000 sequences. This process was repeated 10 times to create 10 test datasets for 500 bp. Corresponding test datasets were also constructed for 1000 bp and 1500 bp. The performance of eukaryotic prediction was evaluated using the F1 score for the eukaryotic class, defined as:

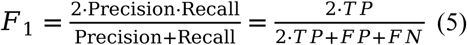

Where TP is True Positives, FP is False Positives, FN is False Negatives. Prediction results are provided in the Supplementary Dataset 2.

#### Viral identification benchmark datasets

The viral identification benchmark utilized a dataset comprising cellular and viral genomes from real metagenomic sequencing across three biomes: gut, seawater, and soil. The original dataset is accessible via the source publication15^15^. Model performance was evaluated using the F1 score (6) for the viral class, Prediction results from different tools are provided in the Supplementary Dataset 3.

### Binning simulation

To simulate metagenomic contig samples with different eukaryotic abundances, we first randomly selected a eukaryotic reference genome from the dataset. This genome was then fragmented into 1500 bp sequences. Based on the number of eukaryotic fragments, bacterial fragments of the same length (1500 bp) were randomly sampled from the entire bacterial dataset according to predefined eukaryotic-to-bacterial sequence ratios. We constructed simulated contig datasets with eukaryotic-to-bacterial sequence ratios of 1:1, 1:2, 1:3, 1:4, and 1:5. For each ratio, the simulation was repeated 10 times, with the eukaryotic sequences kept constant while bacterial sequences were randomly re-sampled each time. For each simulated contigs, MetaBAT2^26^ was employed for binning, and the bin with the highest eukaryotic sequence content was designated as the eukaryotic MAG.

### Eukaryotic MAGs refinement algorithm

According to the MIMAG standards^25^, metagenome-assembled genomes (MAGs) are classified as follows: medium-quality (MQ) MAGs have completeness ≥ 50% and contamination ≤ 10%; high-quality (HQ) MAGs have completeness ≥ 90% and contamination ≤ 5%; all others are classified as low-quality (LQ) MAGs.We adopted these same criteria for eukaryotic MAG quality assessment.

We designed a Eukaryotic MAG Refinement Algorithm to refine eukaryotic LQ MAGs into MQ or HQ MAGs and MQ MAGs into HQ MAGs.The Eukaryotic MAG Refinement Algorithm comprises two main components: searching for an HQ filtering threshold and searching for an MQ filtering threshold. The HQ threshold search process is as follows: First, the Euktect tool is used to calculate the probability of each sequence in the MAG being of eukaryotic origin. A filtering threshold is then set, and sequences with eukaryotic origin probabilities below this threshold are removed. The filtered MAG is evaluated for quality using EukCC^27^. If the MAG meets the HQ standard (completeness ≥ 90%, contamination ≤ 5%), the refined HQ MAG is output. If the HQ standard is not met and the MAG has completeness < 90% and contamination ≥ 5%, the HQ search is terminated. Otherwise, the threshold is adjusted, and the HQ search is repeated. The MQ threshold search follows a similar process: The Euktect tool calculates eukaryotic origin probabilities, a filtering threshold is applied to remove sequences below this threshold, and the filtered MAG is assessed using EukCC. If the MAG meets the MQ standard (completeness ≥ 50%, contamination ≤ 10%), the refined MQ MAG is output. If the MQ standard is not met and the MAG has completeness < 50% and contamination ≥ 10%, the MQ search is terminated. Otherwise, the threshold is adjusted, and the MQ search is repeated. For LQ MAGs, the algorithm first searches for an HQ filtering threshold; if no solution is found, it proceeds to search for an MQ filtering threshold. For MQ MAGs, only the HQ filtering threshold search is performed.

To accelerate the threshold search process in this study, we employed a binary search algorithm. The search range is initialized with a minimum threshold (low = 0) and a maximum threshold high = 1. The midpoint mid = (low + high) / 2 is calculated, and the search range is adjusted based on the evaluation results at this midpoint until a solution is found or the search concludes with no solution.

### Eukaryotic MAGs construction, refinement, and taxonomic labeling

The raw metagenomic sample data and corresponding sample type information used in this study were sourced from previous publication^1^, with specific run accessions detailed in the Supplementary Dataset 4. The contig assembly process was as follows: Raw metagenomic sequencing data were processed using metaWRAP^39^ for quality control, host decontamination, and assembly to generate contigs. Contigs from different samples were binned using MetaBAT2 to produce MAGs.

Since the refinement algorithm refines MAGs by filtering sequences, both completeness and contamination are theoretically non-increasing functions of the filtering threshold. Therefore, input LQ MAGs were required to have completeness ≥ 50%, and input MQ MAGs were required to have completeness ≥ 90%. We applied the Eukaryotic MAG Refinement Algorithm to 407 LQ MAGs and 20 MQ MAGs that met these refinement criteria.

For taxonomic annotation of MAGs, the resulting MQ and HQ MAGs were compared to 5,277 eukaryotic reference genomes using Mash^40^ to calculate the Mash distance (Mash distance = 1 - ANI, where ANI is the average nucleotide identity). For each MAG, the reference genome with the smallest Mash distance was selected as the match reference genome. If the Mash distance between a MAG and its match reference genome was ≤ 0.5, the MAG was considered to belong to the same species as the match reference genome. MAGs with the same match reference genome and a Mash distance ≤ 0.05 (ANI ≥ 95%) were grouped into the same known single genome bin (kSGB). For MAGs with the same match reference genome but a Mash distance > 0.05 (ANI < 95%), Mash distances were calculated internally, and clustering was performed based on a Mash distance ≤ 0.05 to form unknown single genome bins (uSGBs). Visualization of the results was conducted using GraphLAN^41^.Eukaryotic MAGs information is provided in the Supplementary Dataset 5.

### Eukaryotic sequence analysis of food metagenomic samples

The metagenomic samples used for eukaryotic sequence isolation were sourced from the dataset with accession PRJNA997800. Contig assembly for these samples was performed using metaWRAP. The Euktect model was applied to extract eukaryotic sequences from the contigs of these samples. To calculate eukaryotic sequence abundance, extracted eukaryotic sequences were mapped back to the corresponding sequencing data using BWA-MEM^42^. The abundance of eukaryotic sequences was calculated as the ratio of mapped reads to total reads (i.e., eukaryotic sequence abundance = mapped reads / total reads). For cheese-related samples, genomic annotation of eukaryotic sequences was conducted using eggNOG-mapper30. Gene abundance was calculated by mapping gene sequences back to the corresponding sequencing data using BWA-MEM, with gene abundance defined as mapped reads / total reads. Using the annotation results from eggNOG-mapper, gene abundances were regrouped according to their corresponding KEGG Ortholog (KO) identifiers. KEGG pathway enrichment analysis was performed based on KO abundance data using clusterProfiler^43^.

### Eukaryotic sequence analysis of IBD-Related metagenomic samples

The metagenomic samples related to inflammatory bowel disease (IBD) were sourced from the dataset with accession PRJNA400072. Contig assembly was performed using metaWRAP. Contigs smaller than 50 MB were filtered out, resulting in 193 high-quality contig files. The Euktect model was used to extract eukaryotic sequences from these contigs. Sequence abundance for each contig was calculated using BWA-MEM, as described above. Species annotation of the sequences was conducted using Kraken2^8^, with a reference database comprising the 5,277 microeukaryote reference genomes described in the Dataset Preparation section. Species abundance was calculated based on the sequence abundances corresponding to different species-level annotations. The resulting species abundance data are provided in the Supplementary Dataset 6. Random forest models were constructed using Python’s scikit-learn package^44^.

### Fungal multi- phyla model construction and evaluation

To construct the fungal multiclass classification model, we selected data from 10 distinct fungal phyla (*Ascomycota*, *Basidiomycota*, *Blastocladiomycota*, *Chytridiomycota*, *Cryptomycota*, *Microsporidia*, *Mucoromycota*, *Olpidiomycota*, *Sanchytriomycota*, and *Zoopagomycota*) from the superkingdom dataset described in the Dataset Preparation section. Within each phylum, sequences were randomly shuffled and partitioned into training, validation, and test datasets in an 8:1:1 ratio. The model was trained with the following hyperparameters: layer=4, d_model=256, d_inner=1024. Using an input length of 1000 bp, the model was constructed through a pretraining followed by fine-tuning approach, as described in the Training Method section.

To evaluate model performance, test datasets were constructed by randomly sampling 100 sequences of 1000 bp from each of the 10 fungal phyla test datasets, resulting in a total of 1,000 sequences. This process was repeated 10 times to create 10 test datasets for evaluation at 1000 bp. The model’s predictions at 1000 bp were used to compute a confusion matrix. Additional test datasets were constructed for sequence lengths of 2000 bp, 3000 bp, 4000 bp, and 5000 bp, and the F1 score for each fungal phylum was calculated across these datasets.

To investigate the model’s ability to learn sequence features for each fungal phylum, we conducted a sequence clustering analysis. The clustering dataset comprised 100 sequences of 1000 bp from each of the 10 fungal phyla, totaling 1,000 sequences. For the Trinucleotide Frequency (TNF) embedding method, the frequency of each possible 3-mer (trinucleotide, 64 possible combinations) in a DNA sequence was calculated, resulting in a 64-dimensional vector representation. This can be expressed as:

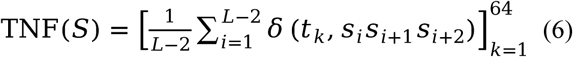

Where S is the DNA sequence, S = s_1_ s_2_…s_L_ *s_i_* ∈ {*A, T, C, G*}, t_k_ is the k-th trinucleotide (64 types in total), δ(*x*, <u>y</u>) is defined as:

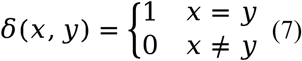

Additionally, the decoder of the fungal multiclass model was removed, leaving only the encoder to embed sequences into vector representations. Clustering was performed on these embeddings using the K-means algorithm from Python’s scikit-learn package. Clustering performance was evaluated using completeness, defined as:

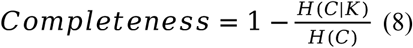

H(C) denotes the entropy of the true class labels, and its calculation formula is

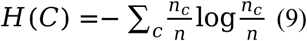

H(C∣K): the conditional entropy of the true labels C given the clustering result K, with the calculation formula as:

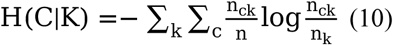

where n_c_ is the number of samples in the true class c, n_k_ is the number of samples in cluster k, n_ck_ is the number of samples that belong to class cand are assigned to cluster k.

The Adjusted Rand Index (ARI) was defined as:

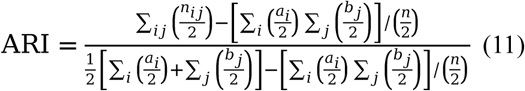

Where n is the total number of samples, n_ij_ denotes the number of samples in the intersection of true class in cluster j, *a_i_* = Σ*_i_ n_ij_* is the number of samples in true class i, *b_j_* = Σ*_i_ n_ij_* is the number of samples in cluster j.

*a_i_* = Σ*_j_ n_ij_* Clustering results were visualized using UMAP^45^.

### Statistical analysis

The statistical significance of differences between the two groups in this study was assessed using the Mann-Whitney U test.

## Data availability

In this study, we used publicly available datasets as detailed in Methods. Metagenomic sample run accessions are listed in Supplementary Dataset 4. Viral benchmark data are available from the https://doi.org/10.5281/zenodo.10886947. The Eukaryotic Identification Benchmark dataset, constructed MQ and HQ eukaryotic MAGs, and additional supporting files can be accessed at https://doi.org/10.6084/m9.figshare.29345837.

## Code availability

The Euktect code and model ckpt&cfg files are available via GitHub at https://github.com/NameFilled/Euktect.

## Author contributions

HL designed the research. YP performed the research. HL and YP analyzed the data. All authors interpreted the results, discussed, drafted and approved the final manuscript.

## Acknowledgements

This work is supported by Shanghai Municipal Science and Technology Major Project, grant 2022YFA0913000 from the National Key R&D Program of China, grant 22208211 and 22378263 from the National Natural Science Foundation of China (NSFC).

## Declaration of generative ai and ai-assisted technologies in the writing process

In preparing this work, the authors utilized ChatGPT-4, Claude Sonnet 4, Google Gemini 2.5 Pro, Grok 3, and Grammarly to enhance writing clarity and ensure consistent formatting. Following the use of these tools, the authors and their research team thoroughly reviewed all content and citations, making necessary edits, and take full responsibility for the final published article.

## Figures and legends

**Extended data Figure 1.**
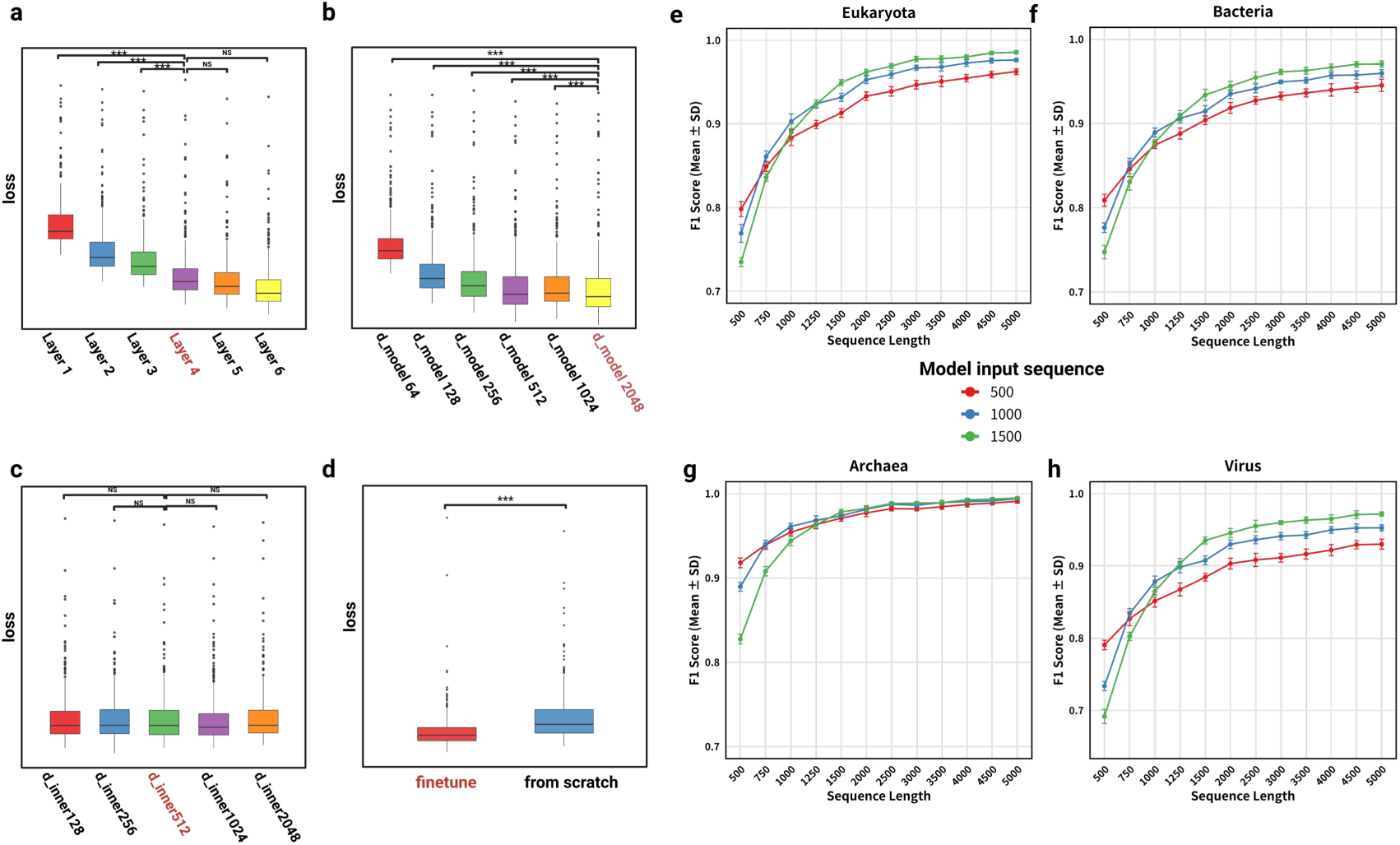
Significance testing of hyperparameters and model performance across superkingdoms on test datasets. Significance between model layer. The final Euktect model was configured with 4 layers. Statistical analysis revealed significant differences in validation loss between the 4-layer model and models with 1, 2, or 3 layers. However, no significant differences were observed when compared to models with 5 or 6 layers, indicating that 4 layers strike an optimal balance for performance (**a**). Significance between Model Width. The final model adopted a width of 2048 units. Significant differences in validation loss were observed between the 2048-unit width and widths of 128, 256, 512, and 1024 units. Due to memory constraints during training, further increases in model width were not explored (**b**). Significance of reverse bottleneck. The model was configured with a reverse bottleneck parameter of 512. Statistical tests showed no significant differences in validation loss between this parameter and other values, suggesting that the reverse bottleneck is not a critical factor in the model’s performance (**c**). Significance between model training methods (**d**). F1 score of eukaryotic sequences across different model and test sequence lengths (**e**). F1 score of bacterial sequences across different model and test sequence lengths (**f**). F1 score of archaeal sequences across different model and test sequence lengths (**g**). F1 Score of viral sequences across different model and test sequence lengths (**h**). ***: *P* value < 0.001, **: *P* value <0.01, *: *P* value < 0.05.

**Extended data Figure 2.**
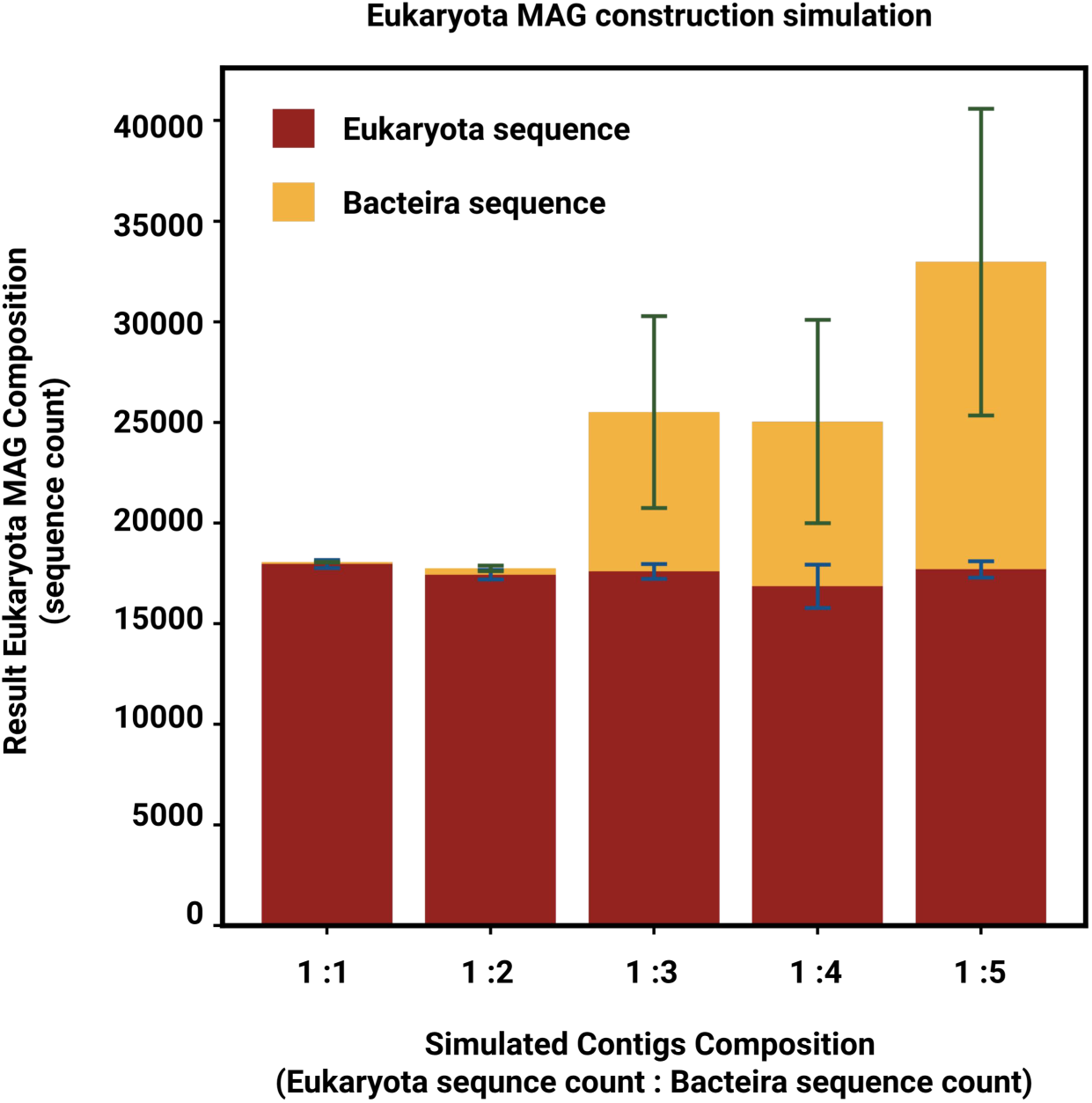
Simulation construction experiment of eukaryotic MAG.

**Extended data Figure 3.**
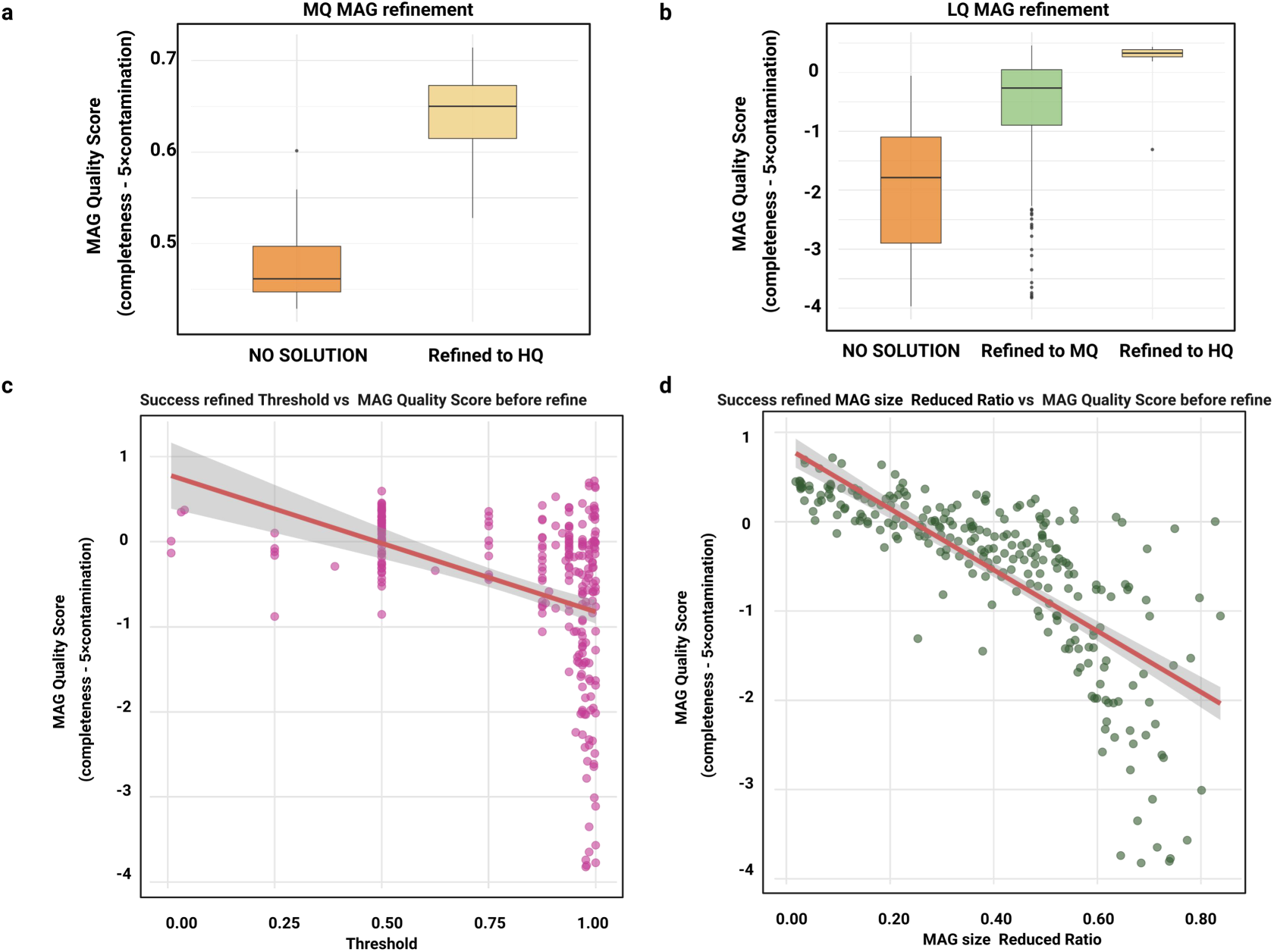
The impact of initial MAG quality on the refinement outcomes. Comparison of original MAG quality scores for successfully refined and No solution LQ MAGs (**a**). Comparison of original MAG quality scores for successfully refined and No solution MQ MAGs (**b**). Relationship between refinement threshold and original MAG quality score for successfully refined MAGs (**c**). Relationship between the MAG size reduction ratio and the original quality score of successfully refined MAGs (**d**).

**Extend data Figure 4.**
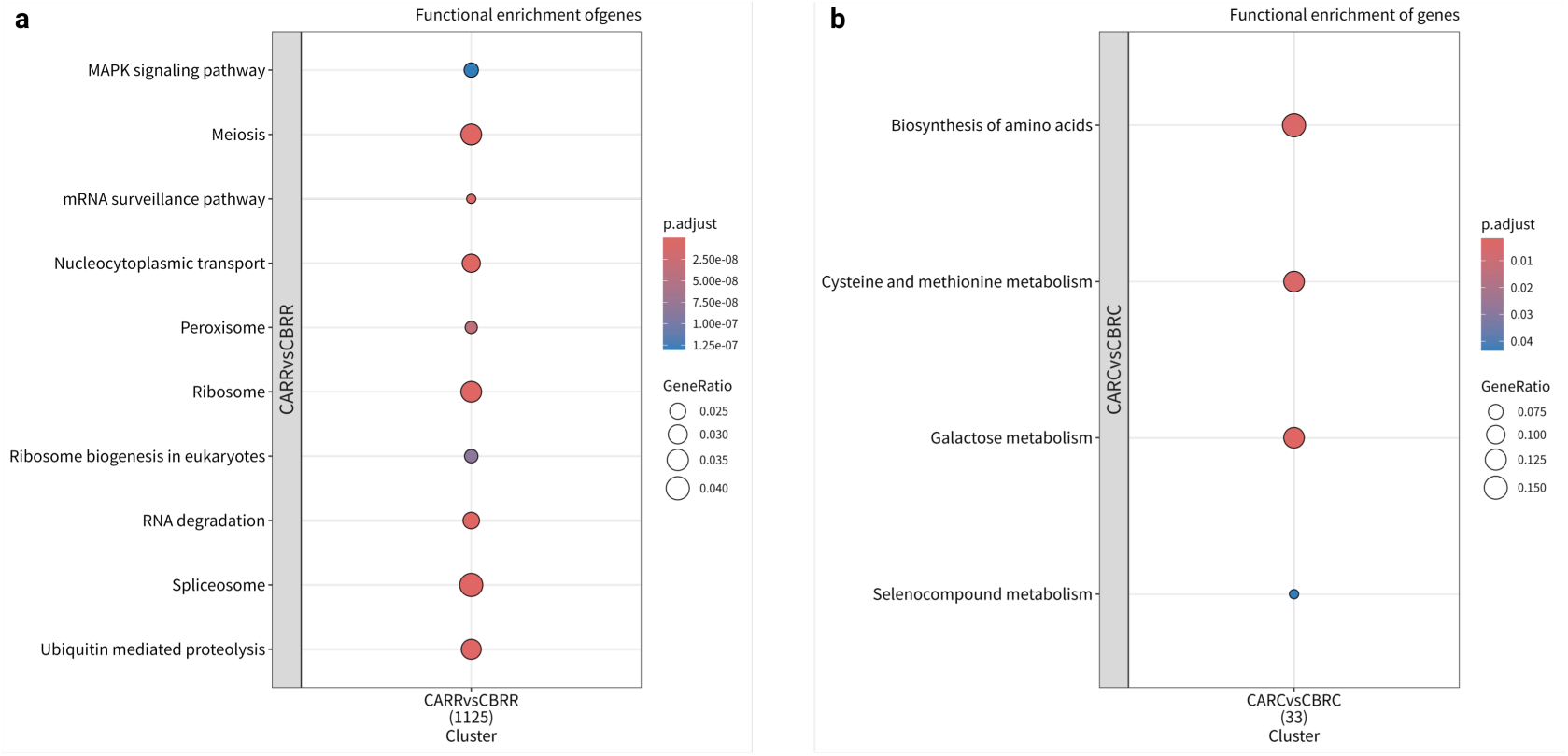
Downstream gene enrichment analysis of eukaryotic sequences. Genes enriched in rind of cheese after ripening samples compared to rind of cheese before ripening samples (**a**). Genes enriched in core of cheese after ripening samples compared to core of cheese before ripening samples (**b**).

**Extended data Figure 5.**
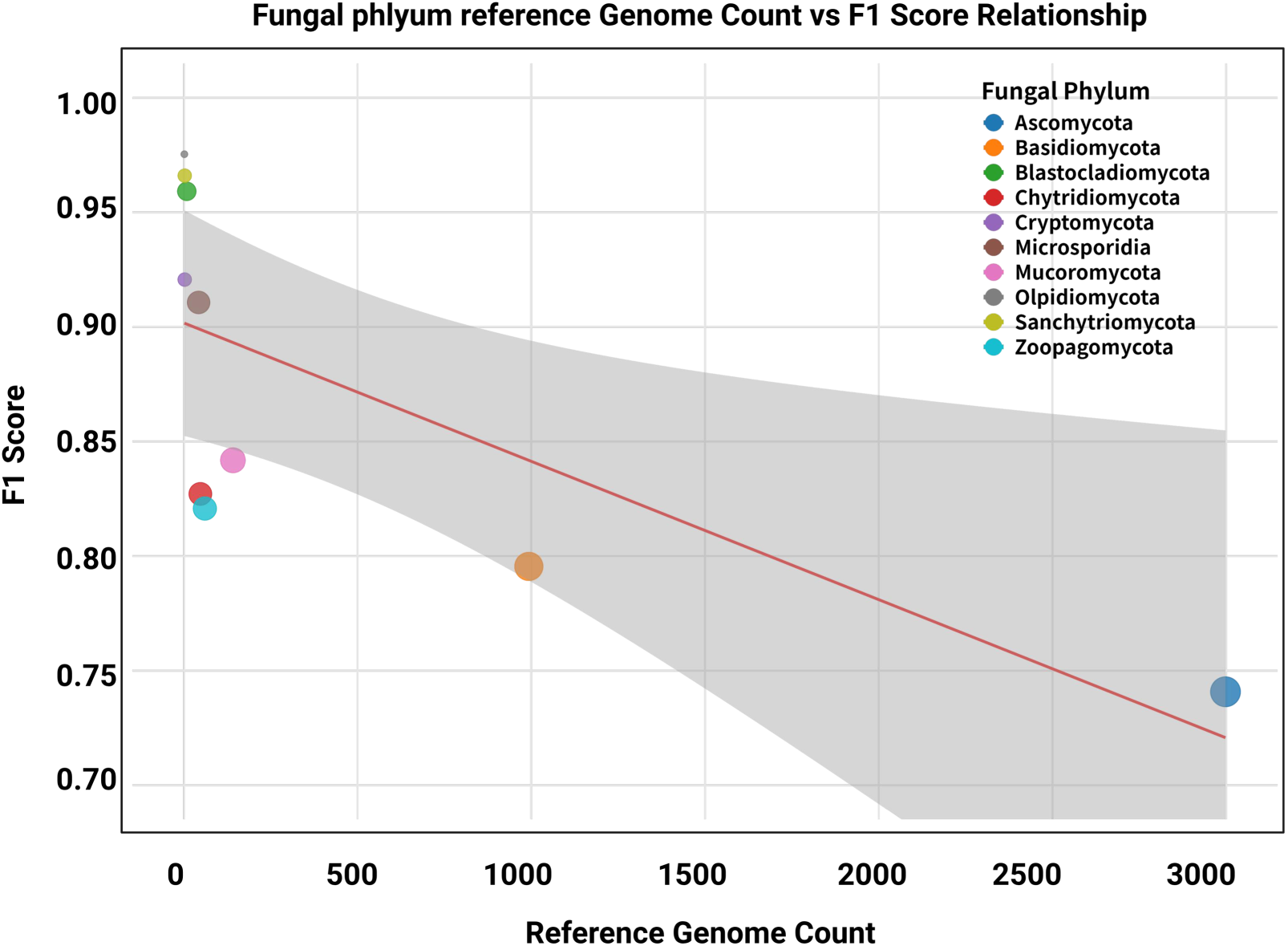
Relationship between the number of reference genomes per fungal phylum and their corresponding F1 scores from the multi-class classification model.

## Supplementary dataset information

Supplementary Dataset 1 dehydrated genome file

Supplementary Dataset 2 Eukaryotic Identification Benchmark Result Supplementary Dataset 3 Virus Identification Benchmark Result Supplementary Dataset 4 Food Sample Run Accession Supplementary Dataset 5 MQ&HQ MAGs information Supplementary Dataset 6 IBD sample eukaryotic species abundance

